# Neuron-Glia Signaling Regulates the Onset of the Antidepressant Response

**DOI:** 10.1101/2021.07.23.453443

**Authors:** Vicky Yao, Ammar Aly, Salina Kalik, Jodi Gresack, Wei Wang, Annie Handler, Anne Schaefer, Olga Troyanskaya, Paul Greengard, Revathy U. Chottekalapanda

## Abstract

Commonly prescribed antidepressants, such as selective serotonin reuptake inhibitors (SSRIs) take weeks to achieve therapeutic benefits^1, 2^. The underlying mechanisms of why antidepressants take weeks or months to reverse depressed mood are not understood. Using a single cell sequencing approach, we analyzed gene expression changes in mice subjected to stress-induced depression and determined their temporal response to antidepressant treatment in the cerebral cortex. We discovered that both glial and neuronal cell populations elicit gene expression changes in response to stress, and that these changes are reversed upon treatment with fluoxetine (Prozac), a widely prescribed selective serotonin reuptake inhibitor (SSRI). Upon reproducing the molecular signaling events regulated by fluoxetine^3^ in a cortical culture system, we found that these transcriptional changes are serotonin-dependent, require reciprocal neuron-glia communication, and involve temporally-specified sequences of autoregulation and cross-regulation between FGF2 and BDNF signaling pathways. Briefly, stimulation of *Fgf2* synthesis and signaling directly regulates *Bdnf* synthesis and secretion cell-non-autonomously requiring neuron-glia interactions, which then activates neuronal BDNF-TrkB signaling to drive longer-term neuronal adaptations^4–6^ leading to improved mood. Our studies highlight temporal and cell type specific mechanisms promoting the onset of the antidepressant response, that we propose could offer novel avenues for mitigating delayed onset of antidepressant therapies.

## Introduction

An estimated 350 million people globally suffer from depressive disorders and these disorders are a leading cause of disability and economic burden worldwide^7^. Selective serotonin reuptake inhibitors (SSRIs) have been used for decades as the first-line pharmacotherapy for depressive disorders due to their safety, efficacy, and tolerability, and are approved for use in both adult and pediatric patients^1, 2, 8, 9^. SSRIs are effective in two-thirds of major depressive disorder (MDD) patients and take weeks before achieving mood improvement^1, 10^. During these weeks, patients can exhibit multiple side effects, among which suicidal ideation is highly predominant^11^. Postmortem and neuroimaging analysis in MDD patients have identified structural and functional impairments within critical neuronal networks during depression pathophysiology^12^. In rodent models, antidepressants are known to raise extracellular serotonin levels within hours of treatment^13^, and are thought to reverse the structural and functional abnormalities by inducing long-term neuronal adaptations such as neurogenesis^14^, altered neuronal morphology and synaptic plasticity^5, 6, 15, 16^. These adaptations, requiring weeks to provide mood improvement in humans and rodent models, indicate the existence of broader, conserved serotonin-dependent mechanisms during response onset which are not understood. Here, we aim to address the delayed cellular and molecular mechanism of action of antidepressants in a mouse depression model using social-isolation rearing stress paradigm and treatment with fluoxetine (Prozac), a widely prescribed SSRI.

### Various cortical cell types respond to stress and antidepressant treatment

To identify the mechanisms that cause weeks-long delay during the antidepressant response, we profiled the temporal dynamics of gene expression changes in response to stress and antidepressant treatment using single-nuclei RNA sequencing (snRNAseq) from the mouse cerebral cortex, an important region in the cortico-limbic mood-circuit^12, 17^ using the 10x genomics platform (**Figure 1A**). Mice were subjected to social-isolation rearing stress to induce depressive-like behavior and were treated with vehicle or fluoxetine as previously described^3, 18^. Gene expression changes for the different groups were analyzed at intervals of early (3 days), week-long (7 days), and close to observed behavioral improvement (10 days) of treatment (**Figure 1B**). Stressed animals displayed anhedonia measured by sucrose preference test (**Figure 1C**) and exhibited heightened startle reactivity as a read-out of their emotional state when their acoustic startle response was measured (**Figure 1D**). To determine the gene expression changes, we first assembled an integrated dataset across all time points and treatment groups. We then identified cell populations based on the corresponding marker genes enriched in each cluster and reviewed cell type specific marker gene expression patterns in available databases (Allen Brain Atlas, GENSAT, gene expression database (GXD) and anatomical studies)^19–25^, and obtained 36 distinct clusters (**Figure 1E**). We grouped and organized these clusters into 15 neuronal (**Extended Figure 1A, Extended Table 1**), 13 non-neuronal (glial and other) **Extended Figure 1B, Extended Table 2**), and 5 inter-neuronal cell types (**Extended Figure 1C, Extended Table 3**).

**Figure 1.**
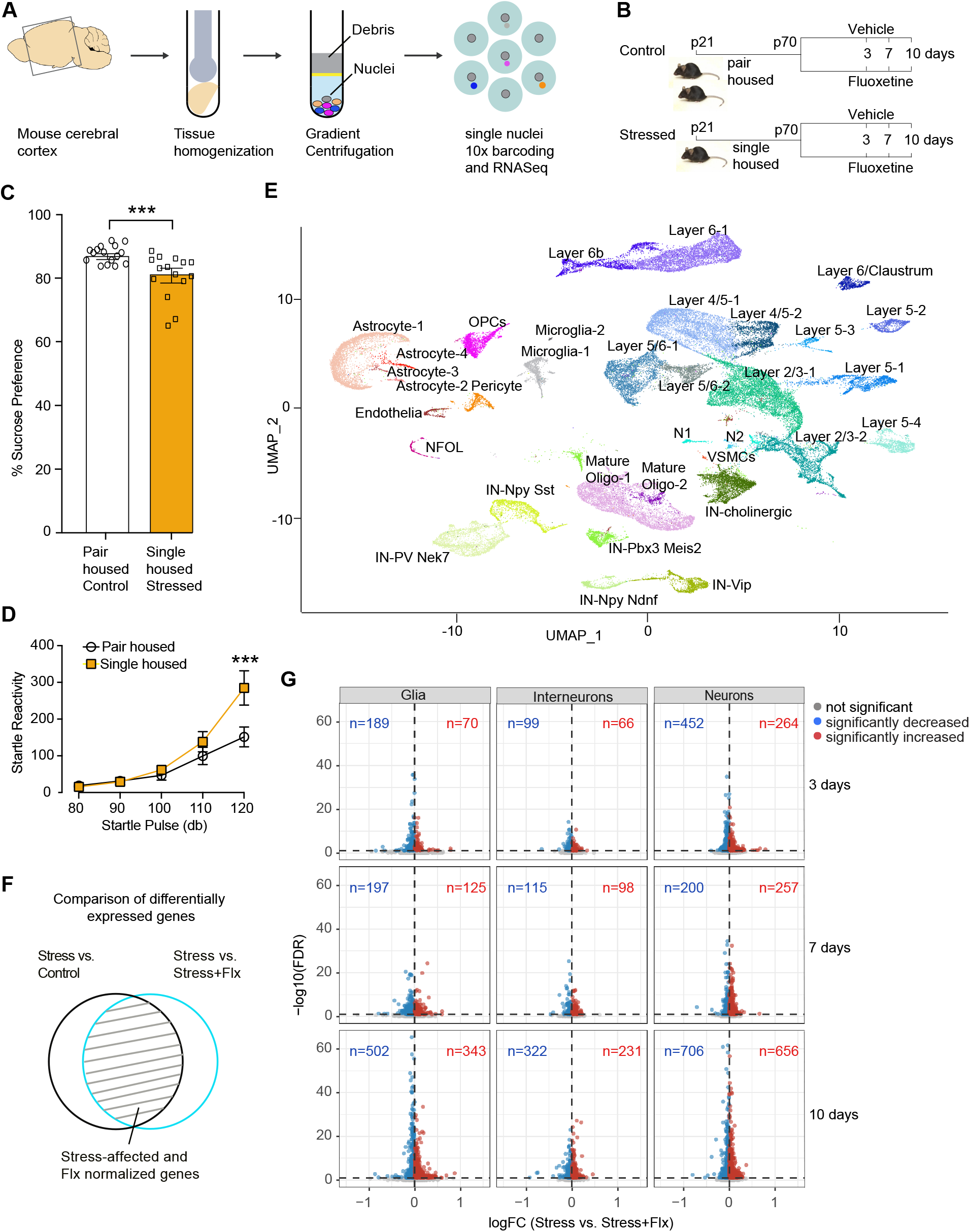
Single nuclei RNA-sequencing analysis of the mouse cortex in response to stress and fluoxetine treatment. **(A)** Mouse cerebral cortex was homogenized, subjected to gradient centrifugation, nuclei purification, single nuclei 10X barcoding, and RNA sequencing. **(B)** Mice were subjected to social isolation rearing stress by single housing from postnatal days P21 until P70, and control mice were pair-housed. Both cohorts were treated with fluoxetine (Flx) for days 3, 7, and 10, n=6 samples each for four treatment groups of Control, ControlFlx, Stress, StressFlx for 3 time points. For each condition, nuclei from n=6 animals were pooled. **(C)** Sucrose preference test (SPT) on control and stressed cohorts to measure anhedonia to assess depressive-like behavior, n=16, pair housed; n=15, single housed on days P64 to P66. **(D)** Acoustic startle response to test reflexive behavior as a readout of emotional state, n=16, pair housed; n=15, single housed on day P68. **(E)** Uniform manifold approximation and projection (UMAP) plot of mouse cortical cells from pooled n=6 samples each of Control, ControlFlx, Stress, StressFlx for 3 time points. Cell cluster labels were annotated based on cell type specific expression of marker genes. Multiple layer subtypes likely representing the same layers from different sub-regions of the cortex were labeled with additional numbers (e.g., Layer 2/3-1 and Layer 2/3-2). Two ungrouped small populations of neurons were called N1 and N2. We identified five inter-neuronal (IN) populations. Vascular smooth muscle cells (VSMCs), two microglial, an endothelial, a pericyte, and four astrocyte populations were identified. Four oligodendrocyte clusters: Oligodendrocyte precursor cells (OPCs); newly formed oligodendrocytes (NFOL); and two populations of mature oligodendrocytes were identified. **(F)** Venn diagram describing the identification of stress-affected Flx-normalized genes by overlaying the differentially expressed genes from the chronically stressed versus control and the chronically stressed versus chronically stressed and treated with fluoxetine comparisons from all cell types. **(G)** Volcano plots showing differential gene expression for stress-affected fluoxetine-normalized genes in the glial, inter-neuronal, and neuronal populations at 3, 7, 10 days (shown for one comparison StressFlx versus stress). Genes significantly upregulated by stress and downregulated by Flx (blue dots), genes downregulated by stress and upregulated by Flx (red dots), and not significant genes (grey dots) are shown. For statistical analysis in **C,** comparisons were made between pair-housed and single-housed mice (n=15-16) using unpaired t-test with Welch’s correction. In **D**, comparisons were made between pair-housed and single-housed mice (n=15-16) that received various intensities of startle pulse using two-way ANOVA and corrections for multiple comparisons was done using Bonferroni’s multiple comparisons test. Data are mean +/− SEM; *P≤ 0.05, **P≤ 0.01, ***P≤ 0.005, ****P≤ 0.001.

To determine the molecular and cellular adaptations triggered by stress that are reversed by fluoxetine treatment, we identified differentially expressed genes that were downregulated by stress and upregulated in response to fluoxetine and genes that were upregulated by stress and downregulated by fluoxetine. We achieved this by overlaying differentially expressed genes from two comparisons: chronically stressed versus control; and chronically stressed versus chronically stressed treated with fluoxetine as depicted in the Venn diagram (**Figure 1F, Extended Table 4**). We found a significant overlap in the transcriptional regulation between the two comparisons in multiple cell types and identified stress-affected fluoxetine-normalized genes. We observed a steady increase in the transcriptional regulation of stress-affected fluoxetine-normalized genes along the time course of treatment. Notably, we found that the normalization of stress-induced changes occurred in cell populations that include glial, neuronal, and inter-neuronal populations, as shown by the volcano plots on day 3, 7, and 10 of treatment (**Figure 1G, Extended Figure 2**). These results highlight the necessity of studying the antidepressant response from the perspective of all the different cell types that work together to shape the behavioral response.

### Neuronal and glial responses to antidepressant treatment are temporally organized

Next, we analyzed the dynamics of the stress-affected fluoxetine-normalized genes at the level of individual glial and neuronal cell populations by performing gene ontology (GO) analysis^26^. We found that both glial and neuronal populations are involved during the early antidepressant response ( **Figure 2A, B and Extended Table 5**). We found significant transcriptional alterations in astrocytes, neurons of layers 2/3, and layer 6 at all time points on days 3, 7, and 10 of fluoxetine treatment. In addition to the above three cell types, oligodendrocyte precursor (OPC) cells also had significant differential expression on day 7 of treatment. Finally, on day 10 of treatment, there were gene expression changes observed in multiple neuronal populations, parvalbumin expressing (PV) interneurons, astrocytes and mature oligodendrocytes (**Figure 2A, B and Extended Table 5**). Changes in mature oligodendrocytes were observed only on day 10 of treatment, concurrent with alterations in multiple neuronal populations. These observations reveal a highly organized response to fluoxetine treatment by the glial and neuronal populations.

**Figure 2.**
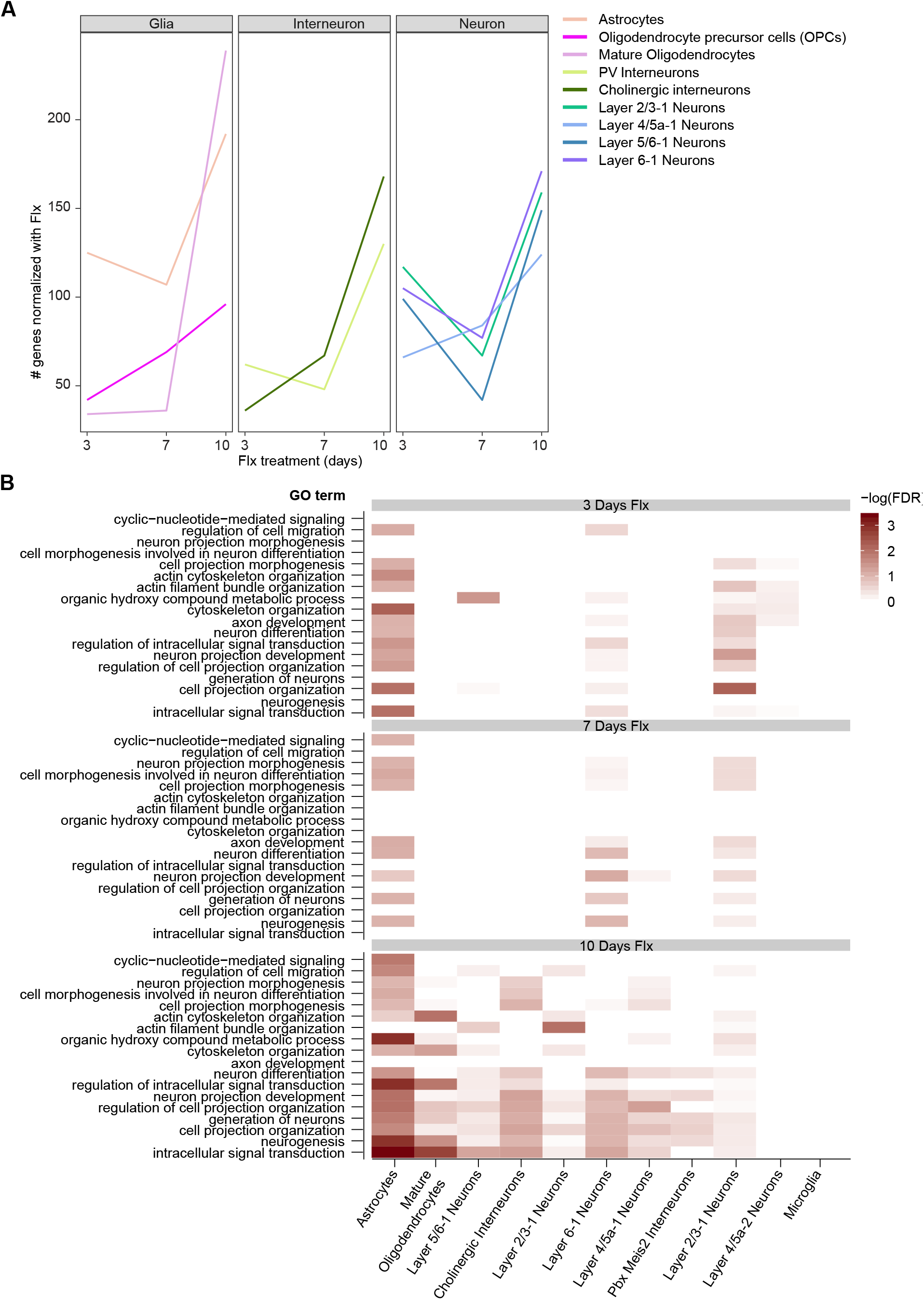
Transcriptional changes in neuronal and glial cell clusters in response to stress and fluoxetine treatment. **(A)** Kinetics of Flx-induced changes in the glial, inter-neuronal, and neuronal cell populations on days 3, 7, and 10 of treatment. Analysis was based on the stress-affected fluoxetine-normalized genes. **(B)** Gene ontology enrichment of the stress-affected fluoxetine-normalized genes on days 3, 7, and 10 of treatment in various cell populations. Analysis based on pooled n=6 samples each of Control, ControlFlx, Stress, and StressFlx at 3, 7, and 10 days of fluoxetine treatment (**Extended Table 3**).

Distinct functions were identified for the stress-affected fluoxetine-normalized gene changes in these cell types as determined by GO analysis and pathway analysis (**Figure 2B)**. Select genes representing cell-type specific cellular functions are represented in **Extended Figure 3 right panel.** Astrocyte changes were observed at all three tested times of treatment. Astrocyte gene changes were attributed to cell projection, cytoskeletal organization, and axonal guidance signaling on days 3, 7 of treatment. In addition, for day 7 of treatment, pathways relating to axonal development, neurotrophin signaling, formation of dendrites, and OPC regulation were observed. Response from multiple neuronal populations were observed on day 10 of treatment, highlighting alterations in neuronal morphology, neurotransmission, and lipid metabolism (**Figure 2B and Extended Figure 3**). Neuronal remodeling and plasticity inducing properties of synaptic long-term potentiation, protein kinase A signaling, synaptogenesis signaling pathway, and cholesterol metabolism were observed. In addition, on day 10 of treatment, mature oligodendrocyte functions contributing to myelination, neuregulin signaling, ephrin receptor signaling, and axonal guidance signaling were observed.

To further characterize the cell type specific functions, we identified upstream regulators modulating stress-affected fluoxetine-regulated genes in glial and neuronal populations. We identified cluster-specific potential regulators such as TCF7L2, RxRB, VEGF, BDNF, and FGF2, molecules that have been previously shown to play a central role in the regulation of stress response in rodent models, depressive-like behavior, and antidepressant responses ^5, 27–30^ (**Extended Figure 3 middle panel**). These molecules have known functions in neuronal plasticity and remodeling pathways, confirming that long-term neuronal adaptations take time to occur. Collectively, these results support the possibility that early changes in the glial cells support late-stage neuronal adaptations contributing to improved behavior.

### Molecular markers to represent the fluoxetine response

Next, we determined whether glial cell functions support late-stage neuronal adaptations that lead to the antidepressant response. First, we asked if the molecular players previously implicated in the antidepressant response had a role in the regulation of neuron-glia interactions. To do so, we chose temporal markers to follow the time course of the fluoxetine response. We had previously identified *S100a10* (p11 protein) as an important molecule which is affected by stress and is essential for providing antidepressant response^3, 31^. Hence we analyzed the dynamics of *S100a10* regulation in response to fluoxetine treatment. Interestingly, we found that *S100a10* transcription is temporally regulated, stimulated only at the late phase, between 9 and 14 days of treatment (**Figure 3A**), closer to the timeline of observed behavioral improvement. This allowed us to use *S100a10* transcription as a proxy for the late response phase and prompted us to identify factors that control *S100a10* transcription to address the cause for the delayed response. p11 is known to be readily induced by growth factors^32^, and therefore, we applied potential growth-factors to stimulate *S100a10* transcription in mouse primary mixed cortical cultures, a model system widely used to study the physiological properties of neurons (**Extended Figure 4 A, B**)^33^. Among tested factors, we observed a strong and rapid induction (within 2h of stimulation) of *S100a10* transcription by BDNF and FGF2 (3-fold) and a small induction by EGF (1.2 fold) (**Figure 3B**). We found that BDNF and FGF2 signaling modulated *S100a10* transcription through the activation of BDNF and FGF2 receptors, TrkB and FGFRs respectively, via regulation of MAPK, PI3K, and JNK cascades (**Extended Figure 4C, D, E, F**). Previously, the significant role of both FGF2- and BDNF-signaling in the regulation of mood, as well as their requirement and sufficiency for providing antidepressant response has been well described^5, 30^. In addition, correlation with respect to reduced levels of both factors was observed in MDD patients and their downregulation in animal models of stress and depression. Thus, our findings and literature precedent led us to choose *Fgf2*, *Bdnf*, and *S100a10* expression as temporal markers to represent the time course of the fluoxetine response.

**Figure 3.**
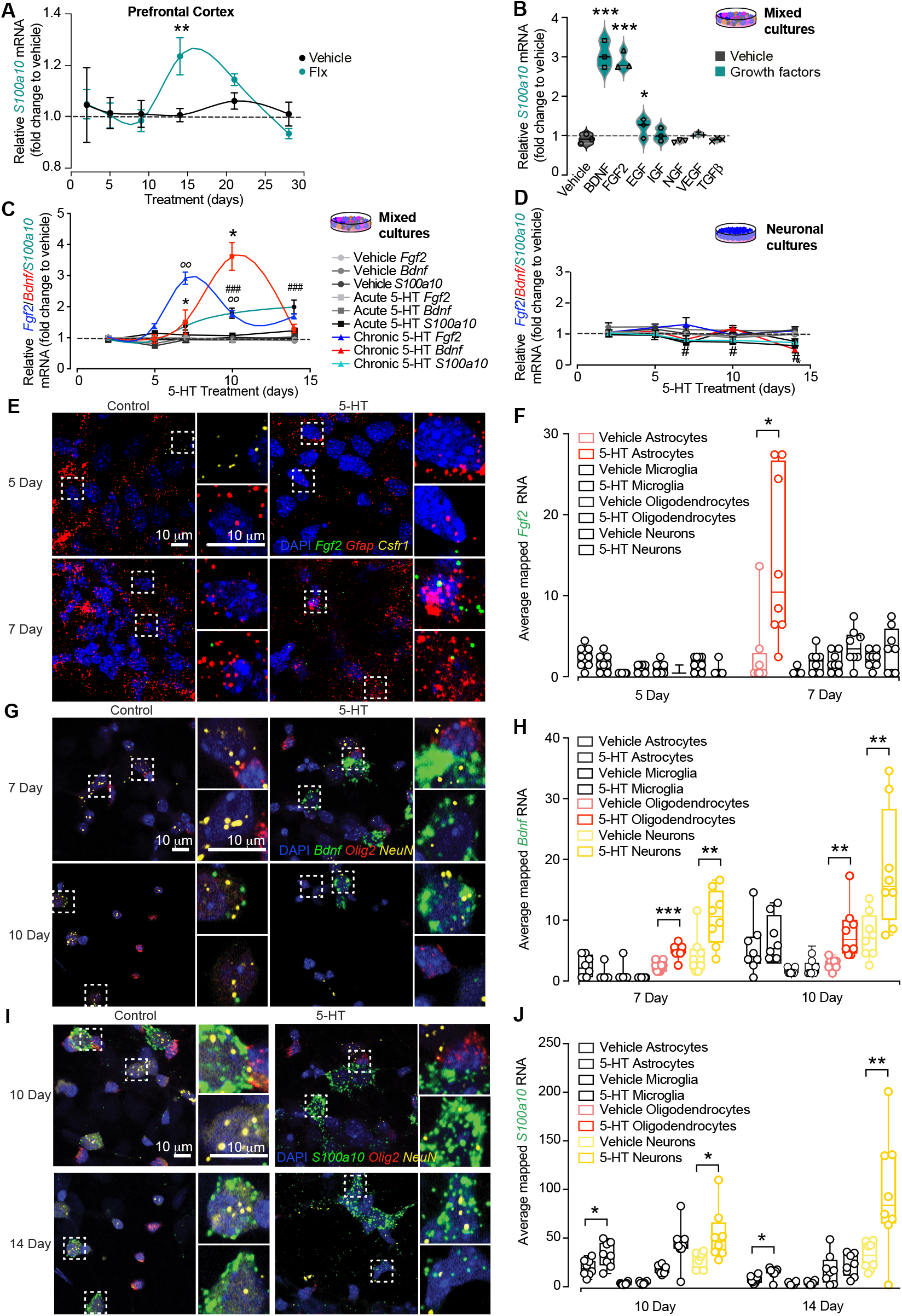
Reproducing the molecular response to fluoxetine in a cortical primary culture system. **(A)** Dynamics of *S100a10* mRNA transcription in the prefrontal cortex (PFC) by quantitative reverse-transcriptase polymerase chain reaction (qRT-PCR) at 2, 5, 7, 14, 21, and 28 days of Flx treatment (n=3). **(B)** Analysis of *S100a10* mRNA transcription in primary cortical mixed cultures by qRT-PCR upon stimulation with growth factors BDNF, FGF2, EGF, IGF, NGF, VEGF, TGFβ for 12h (n=6). **(C)** Analysis of *Fgf2*, *Bdnf,* and *S100a10* mRNA transcription by qRT-PCR at 2, 5, 7, 10, 14 days upon stimulation with vehicle, acute and chronic serotonin (5-HT) treatment in primary cortical mixed cultures (n=6). **(D)** Analysis of *Fgf2*, *Bdnf,* and *S100a10* mRNA transcription by qRT-PCR at 2, 5, 7, 10, 14 days upon stimulation with vehicle, acute and chronic 5-HT treatment in primary cortical neuronal cultures (n=6). **(E)** Fluorescent *in situ* hybridization using RNAscope fluorescent multiplex assay (RNA-ISH) in mixed cortical cultures simultaneously labeled with RNA probes to locate *Fgf2* RNA (green) in astrocytes (using probe for *Gfap* RNA, red) or microglia (*Csfr1* RNA, yellow). **(F)** Quantification of the averaged mapped *Fgf2* RNA in neurons, astrocytes, oligodendrocytes, and microglia. **(G)** RNA-ISH in mixed cortical cultures simultaneously labeled with RNA probes to locate *Bdnf* RNA (green) in neurons (*Rbfox3, NeuN* RNA, yellow) or Oligodendrocytes (*Olig2* RNA, red). **(H)** Quantification of the averaged mapped *Bdnf* RNA in neurons, astrocytes, oligodendrocytes, and microglia. **(I)** RNA-ISH in mixed cortical cultures simultaneously labeled with RNA probes to locate *S100a10* RNA (green) in neurons (*using probe for Rbfox3, NeuN* RNA, yellow) or oligodendrocytes (using *Olig2* RNA, red). **(J)** Quantification of the averaged mapped *S100a10* RNA in the neuronal and glial cell types. For statistical analysis, comparisons were made between vehicle and chronic Flx **(A)** or between vehicle and acute or chronic serotonin (5-HT) treated samples **(C and D),** all (n=3), using two-way ANOVA and corrections for multiple comparisons were performed by running post hoc Tukey’s multiple comparisons test. In **C**, and **D**, p-value significance indicated by symbols **00** for *Fgf2*, * for *Bdnf*, # for *S100a10*. In **B**, comparisons were made between vehicle and growth factor-treated samples (n=3) using one-way ANOVA and corrections for multiple comparisons were done using Bonferroni multiple comparisons test. For statistical analysis in **F, H**, and **J**, comparisons were made between vehicle and chronic 5-HT-treated sections (n=8) for each pair of probes using two-way ANOVA to test effects of 5-HT treatment and time. Corrections for multiple comparisons were performed by running post hoc Tukey’s multiple comparisons test. Data are mean +/− SEM; *P≤ 0.05, **P≤ 0.01, ***P≤ 0.005, ****P≤ 0.001. Experiments were independently reproduced three times with identical results. Scale bar, 10μm.

### Recapitulating the molecular response to fluoxetine in a cortical cell culture system

To address if the chosen molecules essential for the antidepressant response played a role in the regulation of neuron-glia interactions, we asked if we could mimic the molecular response to fluoxetine in a cortical cell culture system. We treated mouse primary mixed cortical cultures harboring both neuronal and glial cell types (**Extended Figure 4A, B**) with serotonin (5-HT), and analyzed the transcriptional regulation of *Fgf2*, *Bdnf*, and *S100a10* expression. Considering that extracellular serotonin levels go up within 30 min of stimulation^13^, we argued that treating cultures directly with fluoxetine would be ineffective, as these cultures lacked the input from the dorsal raphe projection neurons, which provide the main source of serotonin^34, 35^. As expected, we found that fluoxetine treatment of mixed cultures did not change the expression levels of these molecules (**Extended Figure 4G**). Strikingly, we found that chronic treatment of mixed cultures with serotonin stimulated *Fgf2*, *Bdnf*, and *S100a10* transcription when acute treatment had no effects (**Figure 3C, D**). In addition, we observed temporal regulation for *Fgf2* and *Bdnf* transcription, where *Fgf2* stimulation occurred on day 7 and *Bdnf* stimulation occurred between days 7 and 14 of treatment. In parallel sister-cultures comprising predominantly neurons, and where glial cell numbers were largely reduced (**Extended Figure 4A, B**), serotonin treatment failed to stimulate the expression of these genes (**Figure 3C, D**). Thus, with respect to these measured temporal molecular markers, we have reproduced the early *in vivo* molecular response to fluoxetine in a cell culture system, providing evidence that a direct communication between neurons and glial cells is essential. This first approach provided the opportunity to study the properties and interactions between key cell types and molecules that regulate mood.

### Serotonin regulated *Fgf2*, *Bdnf*, and *S100a10* expression within neuronal and glial cell types

To locate which of the glial and neuronal cell types stimulated *Fgf2, Bdnf,* and *S100a10* expression in response to chronic serotonin-treatment, we performed fluorescent *in situ* hybridization detection (FISH) using RNAscope Fluorescent multiplex assay. To detect single molecule RNA targets at the single cell level, we labeled mixed-cultures simultaneously with RNA probes each for *Fgf2, Bdnf, or S100a10* mRNA, and paired them with cell type-specific RNA probes to mark neurons and glial subtypes. We found that chronic serotonin treatment stimulated *Fgf2* mRNA in astrocytes on day 7 (**Figure 3E, F**); *Bdnf* mRNA in neurons between day 10 to day 14 (**Figure 3G, H**); and *S100a10* mRNA in neurons between day 10 to day 14 of serotonin treatment. Notably, *Bdnf* expressing neurons were distinct from those that were expressing *S100a10* (**Figure 3I, J**). The observed effects are strictly temporal and cell type specific. For example, chronic serotonin treatment neither stimulated *Fgf2* mRNA in neurons, oligodendrocytes, nor microglia (**Extended Figure 5A**); nor *Bdnf* mRNA in astrocytes or microglia (**Extended Figure 5B**); nor *S100a10* mRNA in oligodendrocytes or microglia. Low *Bdnf* mRNA expression was observed in oligodendrocytes (**Figure 3G, H**), and low *S100a10* mRNA expression was observed in astrocytes (**Extended Figure 5C**). Additionally, using this system, we also reproduced the temporal dynamics of *Fgf2, S100a10* and *Bdnf* transcription in response to serotonin within neuronal and glial cell types (**Extended Figure 5D, E, F**). Our results demonstrate that the stimulation of key mood-regulating molecules are temporally regulated within an intricate neuron-glia multicellular circuit.

### Neuron-glia secreted factor confers downstream serotonin-dependent function in neurons

Next, we determined if the serotonin-dependent neuron-glia signaling circuit requires physical cell-cell contact or is based on the secretion of neuron-glia factors. To do so, we tested if we could reproduce the serotonin-dependent function by swapping conditioned media from mixed cultures to neuronal cultures during days 5, 7, 10, and 14 of serotonin treatment, and analyzed for *S100a10* transcription (a late-stage temporal marker) (**Figure 4A, B**). Remarkably, we observed a strictly time-dependent induction of *S100a10* in the neuronal cultures that received mixed conditioned media from days 10 and 14 and not from days 5 and 7 of serotonin treatment. We failed to observe induction of *S100a10* when neuronal cultures received vehicle-treated mixed media. These results indicate that serotonin-dependent neuron-glia derived factors secreted from the mixed cultures are temporally regulated and are required and sufficient to stimulate *S100a10* expression in neuronal cultures.

**Figure 4.**
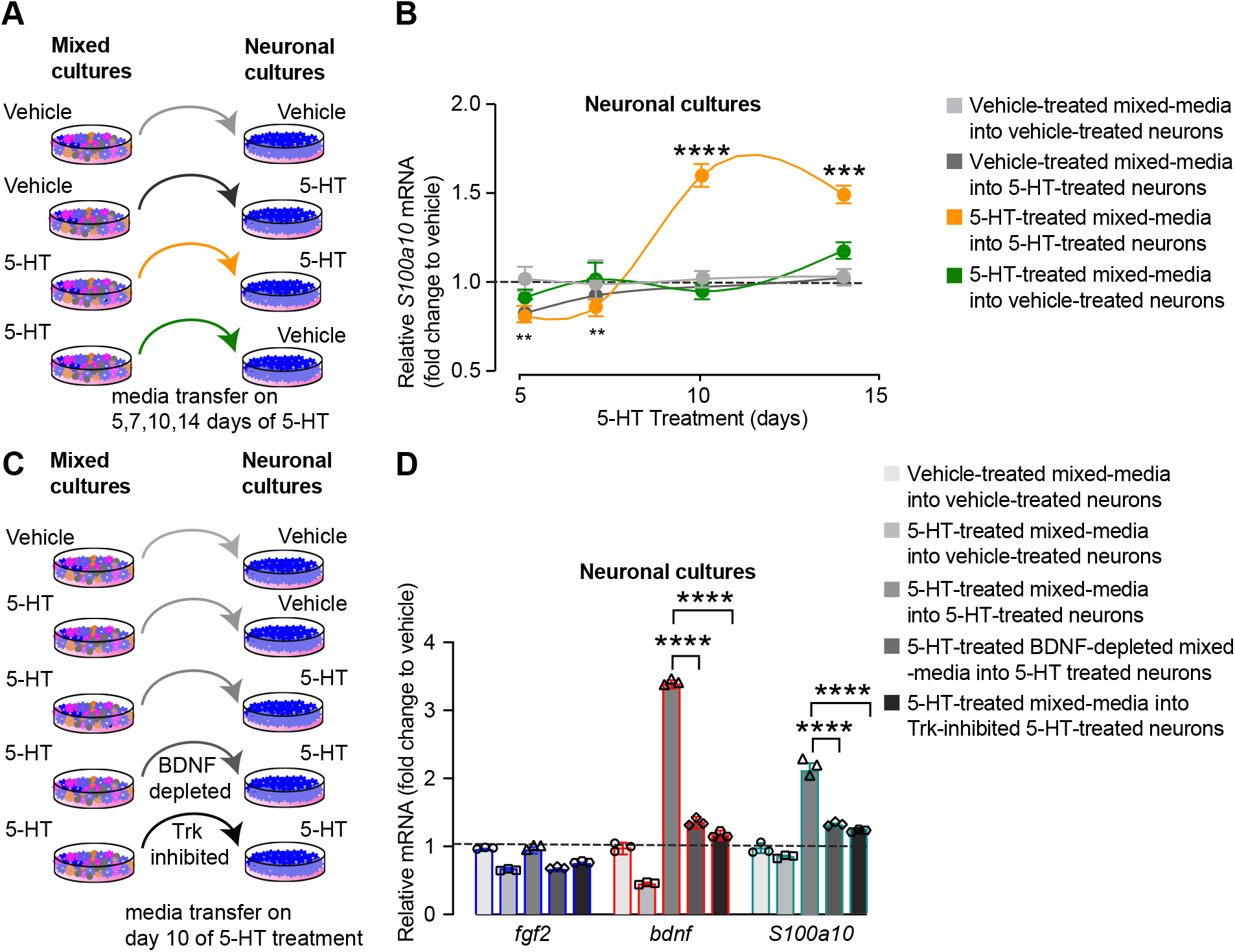
Neuron-glia secreted factor confers downstream serotonin-dependent effects in neurons. **(A)** Schematic showing media transfer from 5, 7, 10, 14 days of vehicle (light grey and black arrows) and chronic 5-HT-treated (orange and green arrows) cortical mixed cultures into 5, 7, 10, 14 days of vehicle (light grey and green circles) and chronic 5-HT-treated (black and orange circles) cortical neuronal cultures. **(B)** Analysis of *S100a10* mRNA expression in cortical neuronal cultures by qRT-PCR after media swap from chronic 5-HT-treated cortical mixed cultures to chronic 5-HT-treated cortical neuronal cultures (n=3). **(C)** Schematic showing media transfer from day 10 of vehicle and chronic 5-HT-treated cortical mixed cultures into day 10 of neuronal cultures of vehicle (two light grey bars), chronic 5-HT-treated (medium grey bar), chronic 5-HT-treated and BDNF-depleted before swap (dark grey bar), and chronic 5-HT-treated Trk autophosphorylation-inhibited (black bar) cortical neuronal cultures. **(D)** Analysis of *Fgf2*, *Bdnf,* and *S100a10* mRNA expression in cortical neuronal cultures after media swap from chronic 5-HT-treated cortical mixed cultures to chronic 5-HT-treated cortical neuronal cultures (n=3). For statistical analysis in **B**, comparisons were made between vehicle and acute or chronic fluoxetine or 5-HT treated samples (n=3) using two-way ANOVA and corrections for multiple comparisons were performed by running post hoc Tukey’s multiple comparisons test. In **D**, comparisons were done using one-way ANOVA, and tested for multiple comparisons using post-hoc Bonferroni multiple comparisons test (n=3). Only comparisons between last three conditions are shown. Data are mean +/− SEM; *P≤ 0.05, **P≤ 0.01, ***P≤ 0.005, ****P≤ 0.001. Experiments were independently reproduced three times.

Our next goal was to identify such neuron-glia derived secretory factors. Best inducible *S100a10* expression was achieved when media was transferred from mixed to neuronal cultures on day 10 of serotonin treatment (**Figure 4A, B**). Also, at that same time point, peak stimulation of *Bdnf* transcription was observed (**Figure 3C**). Therefore, we considered that BDNF could be a potential secreted-factor released from the serotonin-treated mixed cultures that could stimulate *S100a10* expression in neuronal cultures. Hence, we tested this hypothesis using two strategies. We either depleted secreted BDNF from the serotonin-treated mixed-media before transfer using TrkB-Fc to chelate BDNF^36^ or we blocked the TrkB receptor using an inhibitor for Trk receptor autophosphorylation (K252a)^37, 38^ that blocks BDNF/TrkB signaling. Both strategies inhibited the upregulation of *S100a10* in neuronal cultures (**Figure 4C, D**). These results demonstrated that *Bdnf* synthesis and its action on TrkB are both critical steps during serotonin treatment. Additionally, we demonstrated that *Bdnf* synthesis requires neuron-glia interactions and further activation of BDNF-TrkB signaling in neurons is sufficient to upregulate *S100a10* transcription.

### Neuron-glia dependent cross talk between FGF2 and BDNF signaling

To gain further insight into the neuron-glia dependent regulation of *Bdnf* synthesis, we analyzed FGF2- and BDNF-dependent regulation of *S100a10* transcription in mixed and neuronal cultures. We found that FGF2 induced *S100a10* transcription only in mixed cultures, whereas BDNF induced *S100a10* transcription in both mixed and neuronal cultures (**Figure 5A upper panel, Extended figure 6A, B**). These results indicated that upstream FGF2-signaling is likely connected to *Bdnf* synthesis requiring neuron-glia signaling.

**Figure 5.**
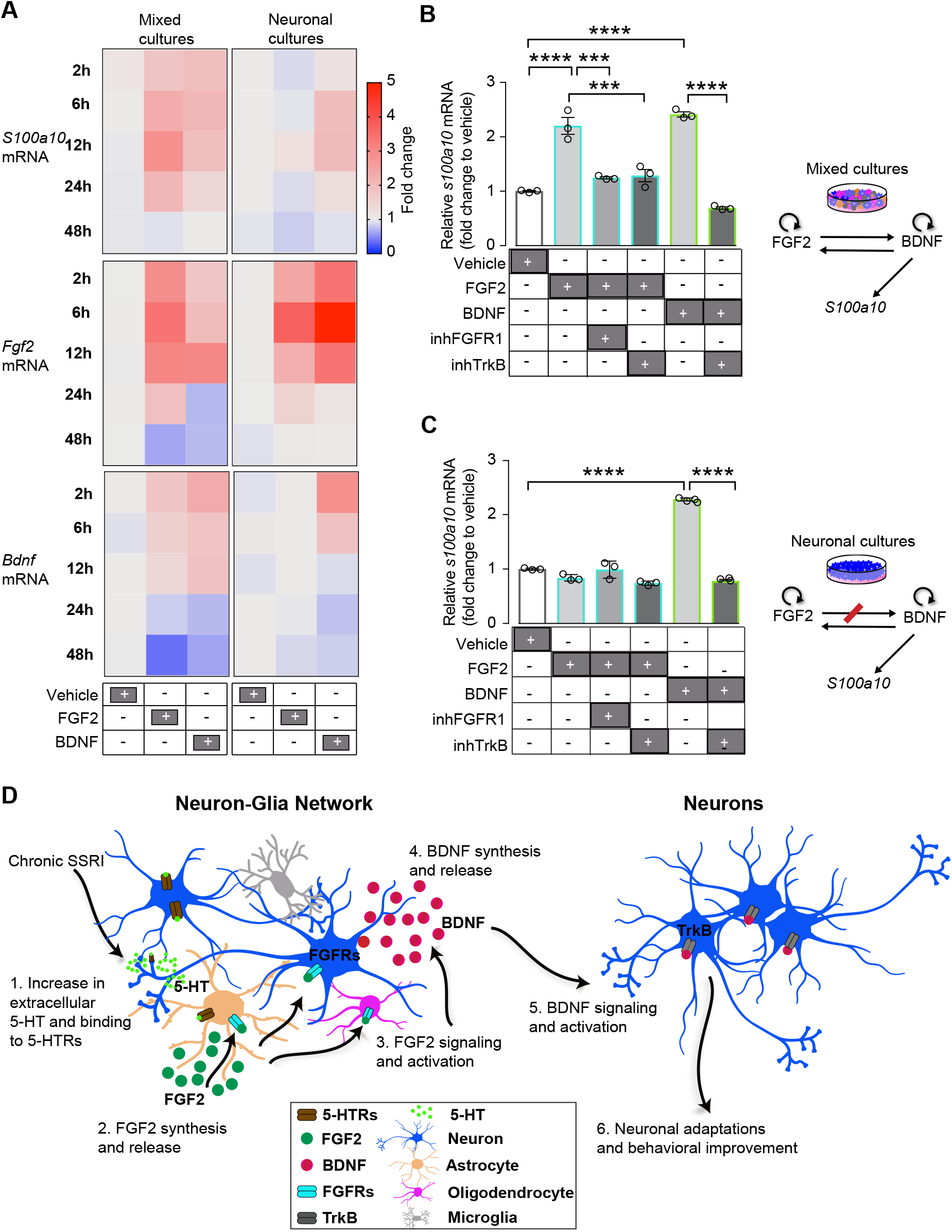
FGF2 regulates *Bdnf* transcription regulating reciprocal neuron-glia interactions. **(A)** Heatmap showing the kinetics of *S100a10, Fgf2*, and *Bdnf* mRNA expression at 2, 6, 12, 24, 48h upon stimulation with vehicle, acute FGF2 and acute BDNF in cortical mixed and cortical neuronal cultures (n=3) based on qRT-PCR analysis shown in **Extended figure 6**. **(B)** Analysis of *S100a10* mRNA expression by qRT-PCR at 8h upon stimulation with vehicle, acute FGF2, acute BDNF, inhibition of FGFR1 (inhFGFR1) 30 min prior to acute FGF2 stimulation, and inhibition of Trk autophosphorylation (inhTrk) 30 min prior to acute BDNF stimulation in cortical mixed cultures (n=3). Schematic showing the interaction between FGF2, BDNF, and *S100a10* in cortical mixed cultures. **(C)** Analysis of *S100a10* mRNA expression by qRT-PCR at 8h of vehicle, acute FGF2, acute BDNF, inhibitor for FGFR1 (inhFGFR1) + acute FGF2 treatment, and inhibitor for TrkB (inhTrkB) + acute FGF2 treatment or acute BDNF treatment in cortical neuronal cultures (n=3). Schematic showing the interaction between FGF2, BDNF, and *S100a10* in cortical neuronal cultures. **(D)** Simplified overview showing how activation of sequential molecular signaling events make up the neuron-glia reciprocal network during chronic serotonin or fluoxetine treatment. For statistical analysis in **A**, comparisons were made between vehicle and acute FGF2 or BDNF-treated samples (n=3) using two-way ANOVA and corrections for multiple comparisons were performed by running post hoc Tukey’s multiple comparisons test. In **B and C**, comparisons were made using one-way ANOVA, and multiple comparisons was tested using post-hoc Bonferroni test. Data are mean +/− SEM; *P≤ 0.05, **P≤ 0.01, ***P≤ 0.005, ****P≤ 0.001. Experiments were independently reproduced three times with identical results.

Next, we analyzed the reciprocal interactions between FGF2 and BDNF-signaling, and observed a strong positive autofeedback regulation. FGF2 application stimulated *Fgf2* mRNA, and BDNF application stimulated *Bdnf* mRNA in both mixed and neuronal cultures (**Figure 5A middle and lower panel, Extended figure 6C, D, E, F**). Intriguingly, we observed a strong reciprocal cross-regulation, where FGF2 stimulated *Bdnf* mRNA only in mixed cultures and not in neuronal cultures. These results demonstrated that FGF2-dependent stimulation of *S100a10* expression occurs via activation of BDNF-TrkB signaling, requiring neuron-glia interactions. Hence, we inhibited TrkB receptor activation and proved that FGF2-dependent stimulation of *S100a10* transcription in mixed cultures occurs via activation of TrkB signaling (**Figure 5B**). As expected, FGF2-stimulation of *S100a10* was downregulated upon inhibition of FGFR1 receptor (predominantly bound to FGF2) and BDNF-stimulation of *S100a10* was downregulated upon inhibition of TrkB receptor activation in both mixed and neuronal cultures (**Figure 5B,C**). These results show evidence that FGF2 stimulates *S100a10* expression only via regulation of *Bdnf* synthesis in a neuron-glia dependent manner, and upon subsequent activation of BDNF-TrkB signaling in neurons (**Figure 5C**).

## Discussion

Patients suffering from MDD take weeks to reverse depressed mood in response to the widely prescribed SSRI class of antidepressants. Interestingly, studies in mouse models indicate a similar timeline to achieve improved mood, suggesting that the molecular and cellular mechanisms underlying delayed response to antidepressants are likely conserved in humans and mice. Our studies provide key insights into the temporal and cell-type specific mechanisms causing the weeks-long delay in producing an antidepressant response. Here we establish that an intricate neuron-glia functional network is recruited by chronic fluoxetine/chronic serotonin treatment to initiate the response. This network involves sequential activation of FGF2-signaling in astrocytes, which directly modulates *Bdnf* synthesis in a neuron-glia dependent manner, and subsequent activation of BDNF-TrkB signaling facilitates neuronal adaptations (via changes in gene expression like *S100a10*), which *in vivo* eventually may lead to improved mood (**Figure 5D)**. We have thus identified a critical missing step in our understanding of the delayed mechanism of action of antidepressants.

We highlight a timely and ordered neuron-glia functional organization regulating the fluoxetine response. Evidence comes from the early and persistent gene expression changes observed in astrocytes, changes in OPCs on day 7, followed by neuronal adaptations occurring in multiple neuronal populations and mature oligodendrocytes on day 10 of treatment. Neuron-glia interactions regulate the development and maintenance of the nervous system^39–41^, and the contributions of glial (astrocytes, oligodendrocytes) and specific neuronal populations in the regulation of mood has been previously reported^12, 16, 42–44^. Ablation of astrocytes is shown to affect depressive-like behavior^45^ and corticolimbic patterns of oligodendrocyte changes are known to occur during MDD^46, 47^. Reproducing the molecular response to antidepressant treatment in a cortical cell-culture system made it feasible to characterize the reciprocal neuron-glia communication and address the precise temporal regulation by FGF2 and BDNF signaling. It is intriguing that extracellular serotonin takes 7 days to trigger *Fgf2* synthesis in astrocytes in the culture system, although we were able to document overall transcriptional changes in astrocytes and layer 2/3 and 6 neuronal populations on day 3 *in vivo*. It is likely that this delay could be due to the serotonin-dependent activation of yet another cell type, whose function leads to the regulation of *Fgf2* synthesis in astrocytes; or simply because the surface regulation of serotonin receptors in neurons or glia require new transcription, translation, or signal transduction events.

Our results confirm that FGF2 signaling and BDNF signaling play a fundamental role in mediating the antidepressant responses ^5, 30^, but we also establish here that glial cells work in concert with neurons to facilitate response onset, subsequently promoting later long-term neuronal adaptations. Conditioned-media swap experiments indicate that BDNF is the neuron-glia derived secreted factor that is sufficient to induce *S100a10* expression (marker for the late phase of the response when behavior could be measured) in neurons. Hence we could argue that it is possible to bypass the neuron-glia interactions and reduce the delay by providing BDNF. Administering BDNF into brains of rodent depression models is known to produce a robust antidepressant response^5^. However, the source of *Bdnf* synthesis is embedded within an intricate neuron-glia interaction. As to the location of the cell type that harbors *Bdnf* synthesis after serotonin treatment, our *in situ* experiments show stimulation of *Bdnf* in neurons. Also, application of BDNF to neuronal cultures stimulates massive *Bdnf* synthesis (**Figure 4D**). It is likely that we missed the exact cell type that synthesizes *Bdnf* or that there are other factor(s) secreted from astrocytes together with FGF2 in response to serotonin, which then facilitates *Bdnf* synthesis in neurons. With prior evidence for *Bdnf* synthesis in other glial cells, such as astrocytes, oligodendrocytes, or microglia^48–50^, it is possible that a small amount of *Bdnf* is synthesized in glial cells, which then stimulates bulk *Bdnf* synthesis predominantly in neurons (**Figure 3G, H**).

Modulation of *Bdnf* synthesis by FGF2 signaling is a critical molecular step, that explains an avenue by which reciprocal neuron-glia interactions are maintained. A prior study documents such a regulation, where FGF2 signaling stimulates the expression of *Bdnf* and of its receptor *TrkB*, post-axotomy of the optic nerve, in order to promote survival of frog retinal ganglion cells^51^. In addition, FGF2 signaling is known to promote astrocyte proliferation^52^, control timely differentiation of oligodendrocytes^53^, and interfere with FGF2 signaling in OPCs to induce depressive-like behavior^54^. These observations, together with the positive autofeedback regulation and cross-regulation between FGF2 and BDNF signaling pathways, indicate that maintenance of multicellular interactions are essential for normal brain function.

Our study provides a greater understanding of the pathophysiology of stress-induced depression and mechanism of action of antidepressants. We establish that neurotrophic pathways of FGF2 and BDNF signaling tightly controls the functioning of an intricate neuron-glia cell circuit to initiate the antidepressant response. We speculate that sustained antidepressant response is possible only upon resetting the disrupted neuron-glia cell circuit by the consequent activation of neurotrophic signaling pathways. Supporting this idea are the observed deficits of reduction in gray matter volume and glial cell density in MDD patients and in animal models of depression^42, 55^; reduced levels of FGF2 and BDNF during stress, depressive-like disorders, and their reversal during antidepressant treatment^5^; and the fact that tissue homeostasis is controlled by growth-factor signaling^56^. Importantly, clinical data indicates that responders to treatment show signs of improvement at day 10 of treatment^11, 57–59^, an early time window when we document adaptations in multiple neuronal populations. We speculate that susceptibility to depressive disorders and treatment resistance could be caused due to dysregulated neuron-glia properties during this early stage. Future investigations to dissect the intricate steps of neuron-glia reciprocal signaling *in vivo* within the cortico-limbic mood circuit could allow for finetuning of the emerging measures to treat mood disorders by modulating neural-circuit dysfunction^60^. Our *in vitro* model system provides a feasible avenue to test for novel antidepressant medications. Further characterizing the critical role for neuron-glia reciprocal interactions during stress and antidepressant response will not only expand our understanding of mechanisms controlling brain homeostasis and repair, but also reveals key features underlying the regulation of complex behaviors.

## Methods

### Animals

Mice were single or pair-housed with a 12h light/dark cycle (lights on from 0700 to 1900 h) at constant temperature (23°C) with *ad libitum* access to food and water. All animal protocols were approved by the Institutional Animal Care and Use Committee (IACUC) at The Rockefeller University.

### Social isolation rearing stress and drug treatment in mice

C57BL/6 background FVB-Tg(*S100a10*-EGFP/Rpl10a) ES691Htz/J mice were subjected to a social-isolation stress paradigm by single housing from P21 until P70 (single housed throughout behavior testing), and the control mice were pair-housed. Both the control and stress cohorts were treated with 0.167 mg/ml fluoxetine hydrochloride (Sigma) in 1% saccharine solution in drinking water as described^18^ for 3, 7, or 10 days. For the PFC fluoxetine experiments, BALB/cJ mice were used as described^3^.

#### Behavior Analysis

##### Sucrose preference test (SPT), a reward based test to measure anhedonia

Mice were presented with two dual bearing sipper tubes in their home cage. One contains plain drinking water and the second contains 2% sucrose solution. Water and sucrose solution intake was measured at 20h with the positions of the bottles switched during the test to preclude side bias. Sucrose preference is calculated as a percentage of the volume of sucrose intake over the total volume of fluid intake averaged over the testing period. Pair-housed mice were temporarily single housed during the course of SPT.

##### Acoustic startle response to measure emotional behavior

The acoustic startle reflex refers to the contraction of the skeletal musculature in response to a high intensity acoustic stimulus. Although the acoustic startle response is a readout for reflexive behavior, this behavior is modulated by emotional state and is an indicator of arousal response. Startle reflex is a physiological measure of emotional regulation. Mice were placed in an animal holder mounted on a startle measuring platform. After an acclimation period, the amplitude of the startle response to acoustic pulse stimuli ranging from 80-120 dB was measured and recorded to determine startle reactivity.

##### Single nuclei purification from the mouse cortex and single cell sequencing using 10X genomics

Mouse cerebral cortex was dissected from 12-week old mice in dissection buffer containing 1x HBSS, 2.5mM Hepes-KOH pH 7.4, 35 mM Glucose, 4 mM NaHCO3. A total of n=24 samples per timepoint comprising six animals per treatment group of Control, ControlFlx, Stress, and StressFlx was collected. Purification of nuclei was achieved as previously described^61, 62^, with modifications. Each cortex was homogenized with a glass Dounce homogenizer (Kimble Chase; 1984-10002) in a lysis buffer containing 20 mM HEPES KOH pH 7.4, 150 mM KCl, 5 mM MgCl2 and the nuclei was pelleted by centrifugation at 1,200xg, 10 min, 4°C. Nuclei was further purified in a 29% iodixanol cushion containing 0.25 M sucrose, 25 mM KCl, 5 mM MgCl_2_, 20 mM tricine pH 7.8 buffer, and myelin floats were removed during each spin. The resulting nuclear pellet was resuspended in a resuspension buffer (0.25 M sucrose, 25 mM KCl, 5 mM MgCl2, 20 mM tricine pH 7.8, EDTA-free protease inhibitor cocktail (Roche, 11836170001)). 6 nuclei samples from each treatment condition was pooled. Nuclei was counted using Countess Cell counter (Thermo Fisher Scientific). 5000 nuclei per sample was processed using 10X genomics using the Chromium Single Cell 3′ Library & Gel Bead Kit v2 (10X Genomics) as per manufacturers protocol. Samples and reagents were prepared and loaded into the chip and droplets were generated using chromium controller for droplet generation. Reverse transcription was conducted in the droplets. cDNA was recovered through de-emulsification and bead purification. Pre-amplified cDNA was further subjected to library preparation and libraries were sequenced on an Illumina Next Seq 500 sequencer high output flow cell as 26 × 57 × 8. Four libraries from each timepoint were pooled together and sequenced on two Next Seq lanes (totally 8 lanes).

##### Single cell Data processing and integration

Seurat’s (v3.1.1)^63^ was used for dimensionality reduction, clustering, visualization, and differential expression analysis for the snRNAseq data. Individual Seurat count matrices was generated from sample output files from CellRanger and subsequently combined into a merged dataset. The percentage of mitochondrial genes were calculated per sample, and per Seurat’s QC recommendations, cells with more than 5% mitochondrial reads, fewer than 500 genes, or more than 5000 identified genes were filtered out. SCTransform was used to normalize and scale the data, including percentage of mitochondrial reads to regress out during the normalization process, and subsequently integrated into a single dataset.

##### Single cell cluster identification and annotation

Seurat’s shared nearest neighbor graph-based Louvain modularity optimization approach was used to cluster the cells. Clustree (v0.4.0) ^64^ was used to choose the appropriate resolution for clustering (0.4). Marker genes consistent for each cluster (across different treatment conditions) were identified via differential expression analysis between cells in each cluster versus all other cells (Wilcoxon rank sum test). Clusters were then assigned based on the identified marker genes. More specifically, cortical neurons were assigned layer-specific identity based on the expression of layer-specific marker genes (layer 2/3, layer 4/5a, layer 5, layer 6 and Claustrum). Multiple layer subtypes were found (they are likely the same layers from different sub-regions of the cortex), and were labeled with additional numbers (e.g., layer 2/3-1 and layer 2/3-2). Two small populations of neurons were ungrouped and hence called N1 and N2. There were 6 inter-neuronal (IN) populations. A small population of vascular smooth muscle cells (VSMCs), two microglial, an endothelial, a pericyte, and 4 astrocyte populations were identified. Four oligodendrocyte clusters were identified: Oligodendrocyte precursor cells (OPCs); newly-formed oligodendrocytes (NFOL); and two populations of mature oligodendrocytes.

##### Identification of fluoxetine-normalized genes and processes

For each cluster and time point (3, 7, and 10 days of chronic fluoxetine treatment), we performed differential gene expression analyses (Wilcoxon rank sum test using Seurat) comparing the following conditions: stress versus control, fluoxetine-treated stress versus stress. Specifically, we aimed to identify genes that were affected by stress, then normalized by fluoxetine. To this end, we identified genes that were significantly differentially expressed in both conditions (FDR < 0.1), and changing in opposite directions (i.e., logFC>0 in stress versus control and logFC<0 in fluoxetine-treated stress versus stress, or vice versa). We then performed Gene Ontology enrichment analysis (for terms in the “Biological Process” aspect of the Gene Ontology) on any cluster and time point that had more than 10 such fluoxetine-normalized genes using goana^26^ from the limma package (v3.38.3).

##### Mouse E18 Primary Embryonic Cortical Cultures

Pregnant C57Bl/6 mice were purchased from Charles River. E18 Embryos were euthanized, and their cortices isolated. Cortices were digested into a single cell suspension using 0.15% Trypsin-EDTA (Gibco, cat # 25200056), followed by mechanical trituration. Cells were then plated in Poly-D-Lysine coated 12-well plates, flasks (Corning, cat # 354470, 356537) at ~2000 cells/mm cell density. Cells were initialing plated in DMEM supplemented with 10% FBS and 1% Penicillin-Streptomycin (Gibco, Cat # 11960069, 10082147, 15070063). For neuronal cultures, plating media was replaced with Neurobasal Media supplemented with B27 and N2 (Thermo Fisher Scientific, Cat # 21103-049, 17504044, A1370701) 3h after initial cell plating. For mixed cultures, half the plating media was replaced as described above, 18h after initial cell plating. For maintenance, half the media was then replaced with Neurobasal media supplemented with B27 and N2 every 3 days. For mixed glia cultures, half the plating media was replaced with plating media 18 hours following initial plating. Half the media was regularly replaced with 10% FBS supplemented DMEM every 3 days. RNA purification from the mixed and the neuronal cortical cultures, that were treated with growth factors FGF2/BDNF and kinase inhibitors; quantitative reverse-transcription and PCR (qRT-PCR), probes for *Bdnf*, *fgf2*, *and S100a10*, and qRT-PCR analysis were all done as described before^3^. For chronic serotonin stimulation experiments, 30 uM of 5-hydroxytryptamine hydrochloride (5-HT) (Millipore Sigma) was applied into DIV5 cortical neuronal and cortical mixed cultures daily, and continued for 14 days. For acute stimulation, serotonin (5-HT) was added only on the day of collection. Cells were harvested 8h after stimulation at each timepoint.

##### RNAscope in Situ Hybridization using RNAscope Multiplex Fluorescent Assay

Mouse primary embryonic mixed cortical cultures were grown in coverslips (NeuVitro, cat #. GG-18-15-PDL) at ~2000 cells/mm cell density. The mixed cultures were treated *Fgf2, Bdnf,* and *S100a10* RNA probes and paired them with cell-type specific RNA probes for neurons and glial subtypes to detect single molecule RNA targets at the single cell level. In situ hybridization was done using RNAscope Multiplex Fluorescent Reagent Kit v2 (ACD Bio, Cat #. 323100) as per manufacturers protocol. Briefly, cells were fixed with 4% PFA in PBS, dehydrated using sequential treatment with 50%, 75%, and 100% ethanol for 5 min, and the coverslips were adhered to superfrost glass slides. The cells were rehydrated using sequential treatment with 100%, 75%, and 50% ethanol for 5 min and permeabilized with 0.1% Tween in PBS. Endogenous peroxidases were blocked using hydrogen peroxide treatment and extracellular proteases were blocked using protease III treatment. RNAscope probes from ACD Bio were hybridized at 40°C. The probes used were as follows: Mm-*Fgf2*, 316851; Mm-*Bdnf*-C4, 424821-C4; Mm-*S100a10*-C2, 410901; Mm-*Rbfox3*-C2, 313311-C2; Mm-*Gfap*-C3, 313211-C3; Mm-*Csfr1*-C2 319141-C2; Mm-Olig2-C3, 447091-C3. Tyramide amplification was performed with the Opal 520, 570, 620, and 690 reagent kits (FP1487001KT, FP1488001KT, FP1495001KT, and FP1497001KT), (Akoya Biosciences) for 1h at 40°C. Coverslips were mounted using ProLong Gold mount and imaged using a Zeiss LSM780 AxioObserver confocal microscope equipped with a Plan-Apochromat 100x/1.4 Oil DIC M27 lens.

Localization of *Bdnf*, *Fgf2*, and *S100a10* RNA in various cell types of neurons, astrocytes, oligodendrocytes and microglia were analyzed. RNA-FISH signal was quantified using the spot counting algorithm as described^65^. Briefly, using ImageJ’s particle analysis feature with a minimum spot size of 0.77 μm, spots were deduplicated along the Z-axis before further processing. Based on the nearest cell identity spot within a radius of 10 μm, each *Bdnf*, *Fgf2*, and *S100a10* RNA signal spot was assigned to one of the four cell types (astrocytes, microglia, neurons, or oligodendrocytes). Signal spots within 10 μm of multiple cell identity spots were excluded unless its nearest neighbor was at least 5 μm closer than its next nearest neighbor. ~40% of all detected spots were assigned to one of the four cell types with the described criteria. Eight representative images for the localization of each gene per time point was quantitated.

Expression levels of *Fgf2*, *S100a10*, and *Bdnf* RNA in response to serotonin treatment were quantitated as follows. RNA expression spots were counted using ImageJ particle analysis feature and were assigned to the cell types as described as above. We had samples from 4 time points for *Bdnf* (days 5, 7, 10, 14 of 5-HT treatment), and 2 points each for *Fgf2* and *S100a10* (days 10 and 14 of 5-HT treatment) respectively. For each time point, the number of RNA spots of *Bdnf*, *Fgf2*, and *S100a10* RNA in each image was counted and 8 images per cell type specific marker set was analyzed. Set 1 labeled by *Csfr1* (microglia) and *Gfap* (astrocytes) and set 2 labeled by NeuN (neurons) and *Olig2* (oligodendrocytes) and the spots are assigned to a cell type. Average RNA/cell type was then computed from the data. Hence a total of 16 data points per timepoint for each of the *Bdnf*, *Fgf2*, and *S100a10* RNA datasets were analyzed.

##### Quantification and Statistical Analysis

Graphs and statistical analysis was done using GraphPad Prism 9. Statistical details is included in the figure legend for each experiment. Briefly, for two group comparisons, we used two-tailed unpaired student’s t-test. For multiple group comparisons, we used one-way or two-way ANOVAs and corrections were applied using the appropriate post hoc test. In all experiments, p value < 0.05 was considered significant. Bar graphs show mean values and the error bars are standard error of the mean (± SEM). The number of animals for each experiment and biological replicates for cell culture experiments are denoted by n. For behavior experiments, all studies were carried out and analyzed with the experimenter blinded to the treatment group.

## Acknowledgements

We thank Drs. Marc Flajolet, Priya Rajasethupathy, Dipon D.Ghosh, Jean-Pierre Roussarie, and Ms. Debra Poulter for critical comments on the manuscript. We thank Elisabeth Griggs for assistance with graphics. This research was supported by The JPB Foundation (to P.G), NARSAD Young Investigator Grant from the Brain & Behavior Research Foundation (to R.U.C), NIH R01 GM071966 (to O.G.T), V.Y. was supported in part by US NIH grant T32 HG003284 (to O.G.T). O.G.T. is a senior fellow of the Genetic Networks program of the Canadian Institute for Advanced Research (CIFAR).

## Author contributions

R.U.C., V.Y., A.S., O.G.T., and P.G. designed and planned the experiments. R.U.C., V.Y., and O.G.T. planned the single cell experiments. V.Y., and W.W. performed the single-cell data-analysis and interpretation. R.U.C., J.G., A.A., A.H. conducted drug treatments and mouse behavioral testing. A.A., and S.K. genotyped the mice. R.U.C., S.K., and A.A. planned and performed cell culture, serotonin treatment, inhibitor treatments, qRT-PCR and analysis. R.U.C., and A.H. performed western blots and analysis. R.U.C., and A.A. planned and performed RNAscope experiments. All authors contributed to the writing of the manuscript.

---

**Extended Figure 1.**
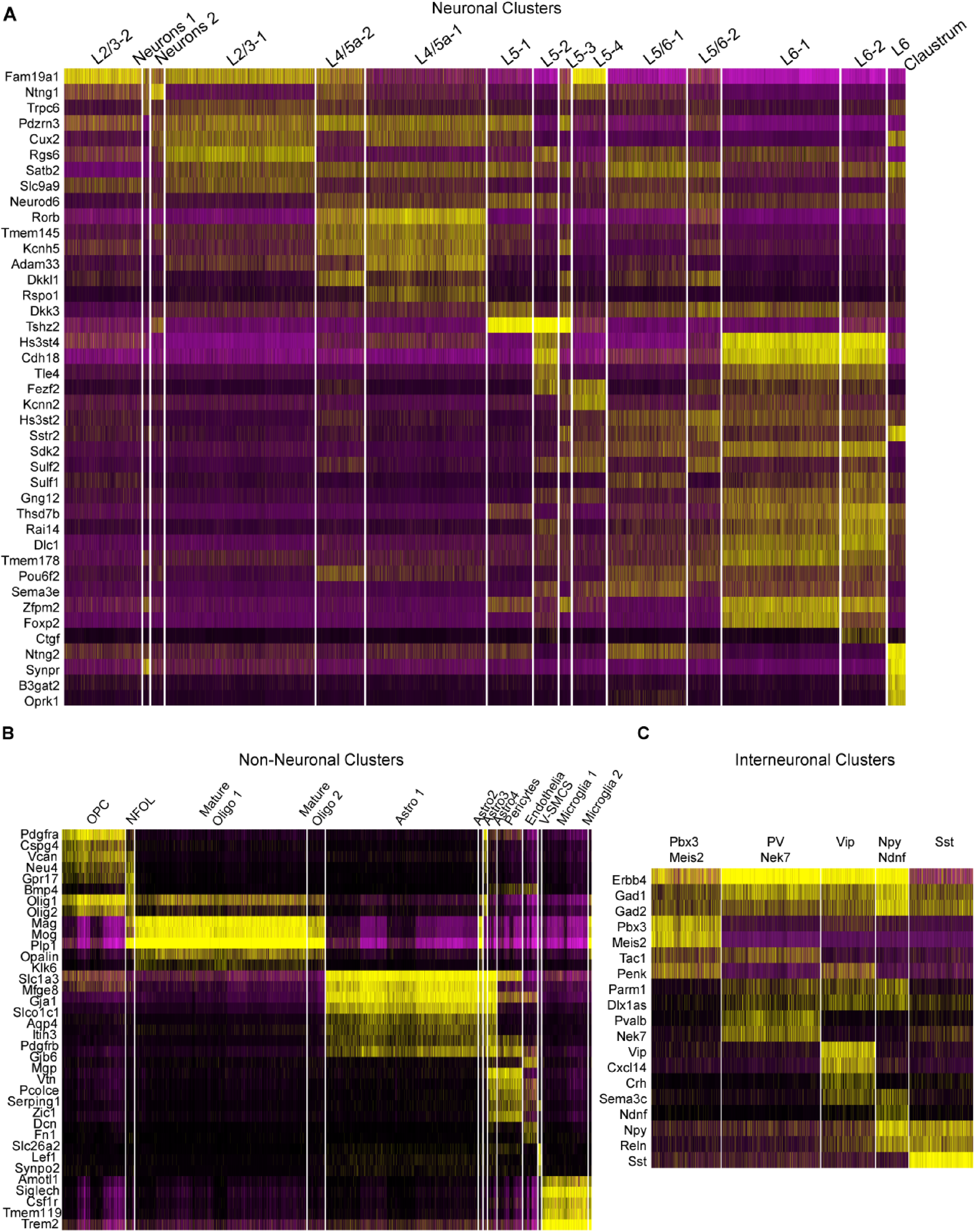
Classification of neuronal, non-neuronal and inter-neuronal cell type clusters based on cell type specific marker gene expression. **(A)** Heatmap showing the expression of cell-type specific marker genes in the cortical neuronal clusters. Cortical neurons are assigned layer-specific identity based on the expression of layer-specific marker genes. Each row represents the expression of a single gene, and each column represents a single nuclei from a single cell, and many such cells with similar gene expression profiles are clustered and grouped. **(B)** Heatmap showing the expression of cell type specific marker genes in the cortical non-neuronal clusters. **(C)** Heatmap showing the expression of cell type specific marker genes in the inter-neuronal clusters. Samples are n=12 comprising four treatment groups of control, controlFlx, stress, stressFlx at 3 time points. For each time point, for each treatment group, nuclei from n=6 animals were pooled.

**Extended Figure 2.**
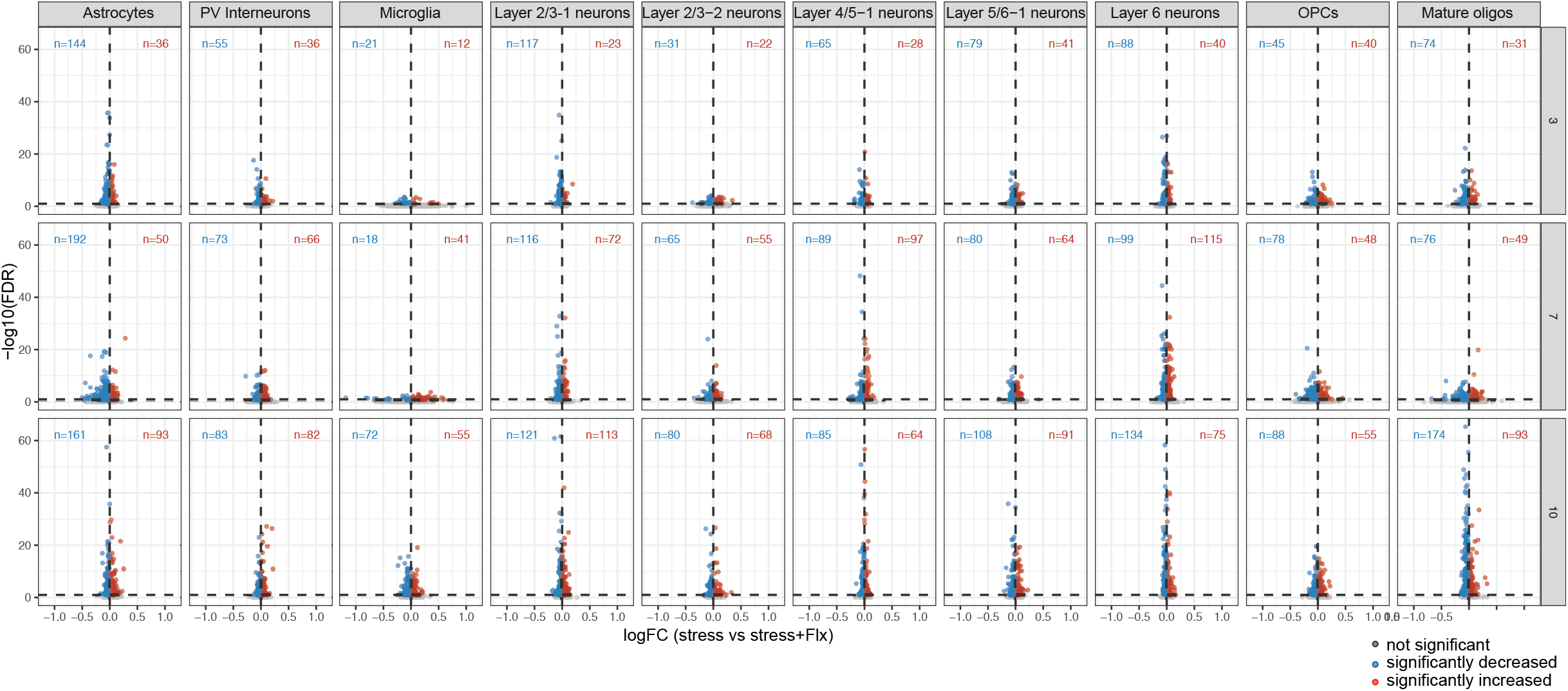
Volcano plots showing the differential expression of stress-affected fluoxetine-normalized genes. Differential gene expression for stress-affected fluoxetine-normalized genes in the glial, inter-neuronal, and neuronal populations at 3, 7, 10 days (shown for one comparison of Stress versus StressFlx). Genes significantly upregulated by stress and downregulated by Flx (blue dots), genes downregulated by stress and upregulated by Flx (red dots), and not significant genes (grey dots) are shown. Genes that are significantly differentially expressed in both conditions (FDR < 0.1), and changing in opposite directions (logFC>0) in stress versus control and (logFC<0) in fluoxetine-treated stress versus stress, or vice versa).

**Extended Figure 3.**
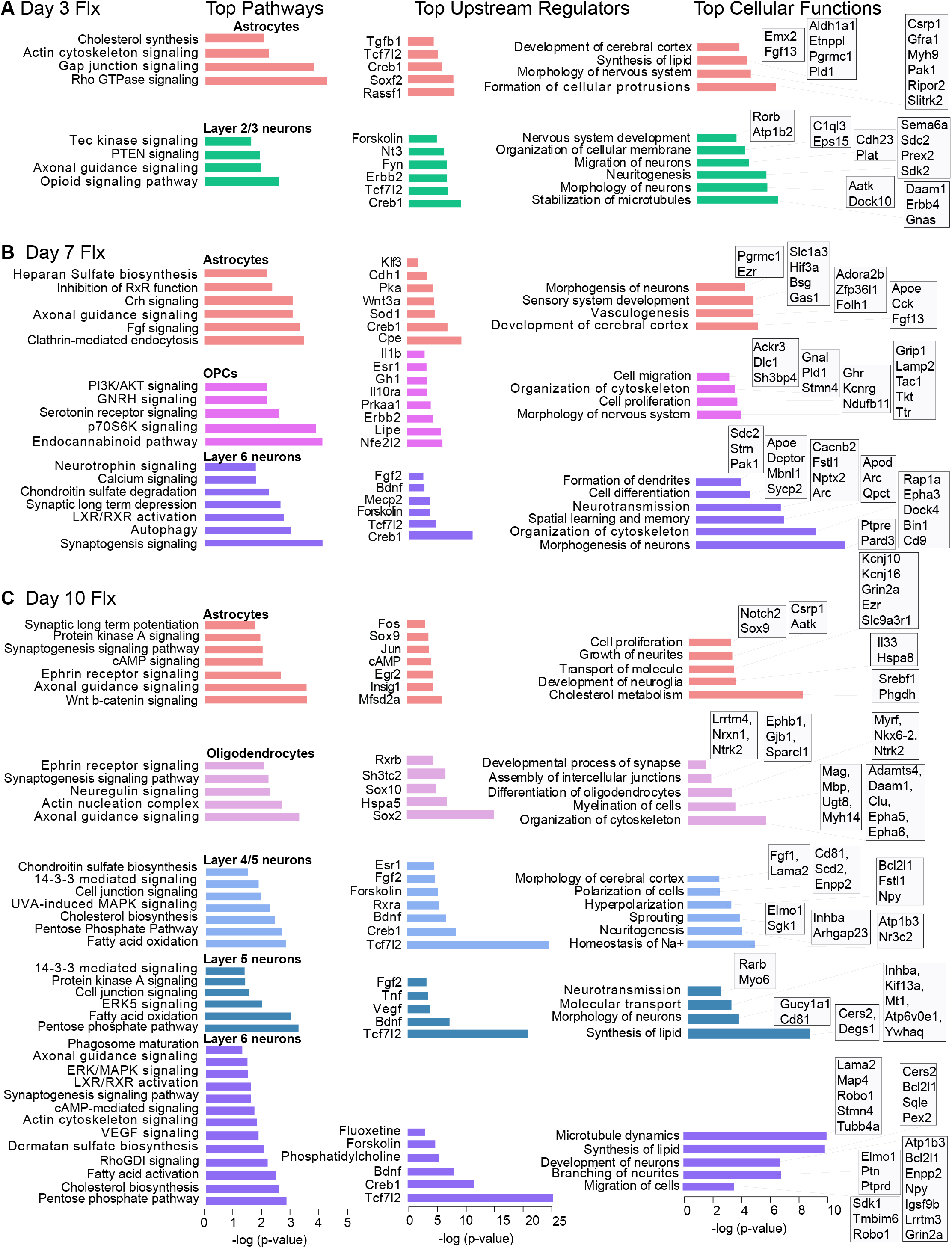
Cell type specific molecular and cellular functions regulated by the stress-affected fluoxetine-normalized genes by Ingenuity Pathway Analysis. **(A)** Enriched pathways, top upstream regulators, and enriched cellular functions in astrocytes and Layer 2/3 neurons on day 3 of Flx treatment. **(B)** Enriched pathways, top upstream regulators, and enriched cellular functions in astrocytes, OPCs, and Layer 6 neurons on day 7 of Flx treatment. (**C)** Enriched pathways, top upstream regulators, and enriched cellular functions in astrocytes, mature oligodendrocytes, Layer 4/5a neurons, Layer 5/6 and Layer 6 neurons on day 10 of Flx treatment. The significant values (p-value of overlap) for enriched pathways and enriched cellular functions indicate the probability of association of molecules from the dataset with the canonical pathway by random chance alone and is calculated by the right-tailed Fischer’s exact test and is displayed along the x-axis. Upstream regulator analysis identifies molecules that affect the expression, transcription or phosphorylation of genes in the experimental dataset, and the significant overlap p-value (P< 0.05) is based on right-tailed Fisher’s exact test.

**Extended Figure 4.**
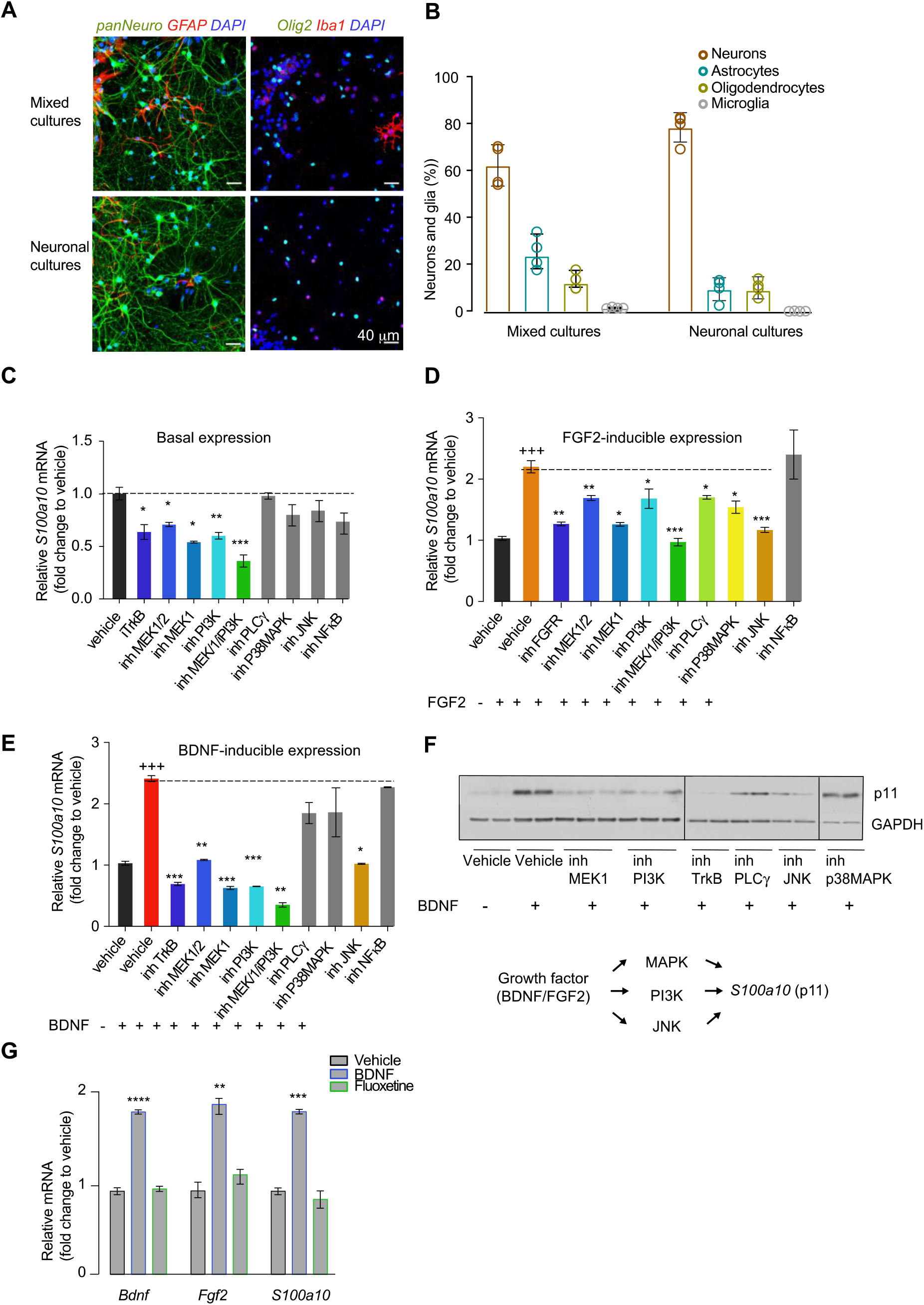
Primary cortical culture cell types and stimulation of *S100a10* mRNA by growth factors and fluoxetine. (**A**) Representative images of immunofluorescence analysis in DIV12 (days in vitro) primary cortical mixed and neuronal cultures to label neurons (pan-neuronal antibody), astrocytes (GFAP antibody), oligodendrocytes (Olig2 antibody), microglia (Iba1 antibody), and all nuclei (DAPI). (**B**) Cell counting of neuronal and glial cell types using ImageJ analysis. Cells were assigned to one of the four cell types based on the intensity of fluorescent signal from each cell marker staining. Cells were counted from n=3 sections, n=3 replicates and classified per data point. (**C**) basal (**D**), FGF2- (**E**), and BDNF-inducible *S100a10* mRNA expression upon pharmacological inhibition of the receptor tyrosine kinase pathways in cortical mixed cultures. Inhibitors for TrkB (K252a), FGFRs (PD 173074), MAPK (U0126, MEK1/2 inhibitor), PI3K (LY294002), PLC_γ_ (U73122), p38MAPK (SB203850), JNK (SP600125), NFkB (SN50) were used. Figure legends have an “inh” for inhibitors. (**F)** Western blots showing the effect of kinase inhibition on BDNF-induced expression of p11 protein (*S100a10 gene*) in cortical mixed cultures. GAPDH was used as a loading control. Also shown in **F** is a schematic diagram summarizing the activation of FGF2 and BDNF-inducible kinase cascades that lead to *S100a10* regulation. (**G**) qRT-PCR analysis of *Bdnf, Fgf2* and *S100a10* mRNA expression at 12h after acute treatment with BDNF (purple bar) and Flx (green bar) in cortical mixed cultures. Statistical analysis in **C**, **D**, **E**, **G** was done using one-way ANOVA and corrections for multiple comparisons were performed using post-hoc Bonferroni test. Comparisons were made between growth factor-treated samples with and without inhibitors in **C, D and E**; n=6. Fold change values for the control vehicle samples are normalized to 1. Comparisons were made between vehicle- and Flx-treated samples in **G;** n=3. In **D** and **E**, + symbol indicates comparison between vehicle- and BDNF- or FGF-treated, and * symbol indicates comparison between growth factor-treated samples with and without inhibitors. Data are mean +/− SEM; *P≤ 0.05, **P≤ 0.01, ***P≤ 0.005. Dashed lines in **C** indicate fold change of the control sample and dashed lines in **D** and **E** indicate fold change of the FGF2- and BDNF-treated sample. Experiments in **C**, **D**, **E**, **G** were reproduced three times with identical results.

**Extended Figure 5.**
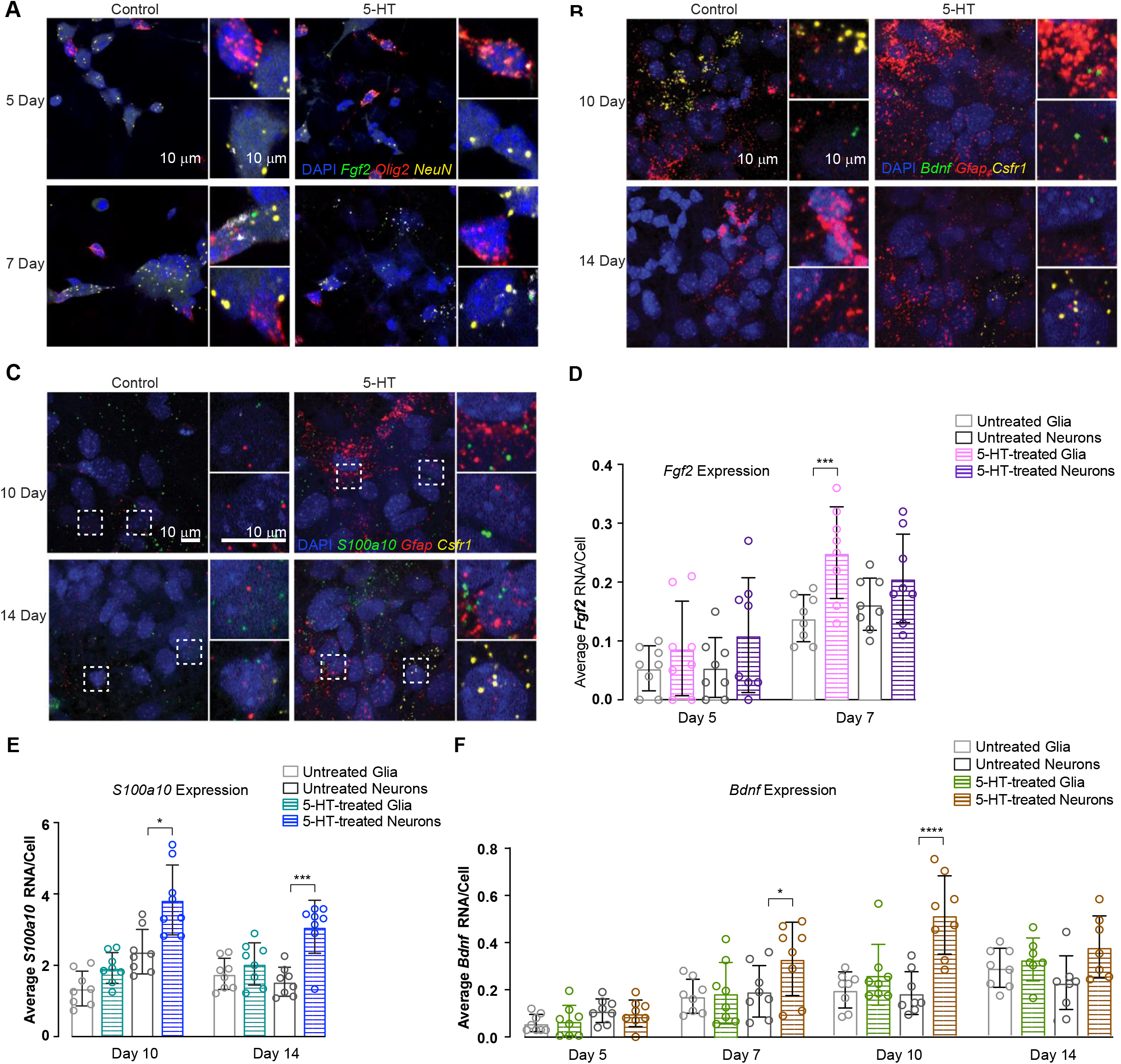
RNAscope fluorescent *in situ* hybridization multiplex assay (RNA-ISH) in cortical mixed cultures to analyze expression of *Fgf2, Bdnf, and S100a10* RNA in neuronal and glial cell types. **(A)** Representative images of RNA-ISH in mixed cortical cultures simultaneously labeled with RNA probes to locate *Fgf2* RNA (green) in oligodendrocytes (*using probe for Olig2* RNA, red) and neurons (using *Rbfox3/NeuN* RNA, yellow). **(B)** Representative images of RNA-ISH in mixed cortical cultures simultaneously labeled with RNA probes to locate *Bdnf* RNA (green) in astrocytes (using probe for *Gfap* RNA, red) and microglia (*using probe for Csfr1*, yellow). **(C)** Representative images of RNA-ISH in mixed cortical cultures simultaneously labeled with RNA probes to locate *S100a10* RNA (green) in astrocytes (using probe for *Gfap* RNA, red) and microglia (*using probe for Csfr1*, yellow). Experiments were reproduced with identical results. **(D)** Quantitation of *Fgf2* mRNA between untreated and 5-HT-treated neurons and glia. **(E)** Quantitation of *S100a10* mRNA between untreated and 5-HT-treated neurons and glia. **(F)** Quantitation of *Bdnf* mRNA between untreated and 5-HT-treated neurons and glia. For statistical analysis in **D, E**, and **F**, comparisons were made between untreated and chronic 5-HT-treated sections (n=8) for each pair of probes using two-way ANOVA to test the effects of chronic 5-HT treatment and time. Corrections for multiple comparisons were performed by running post hoc Tukey’s multiple comparisons test. Data are mean +/− SEM; *P≤ 0.05, **P≤ 0.01, ***P≤ 0.005, ****P≤ 0.001. Experiments were independently reproduced three times with identical results. Scale bar, 10μm.

**Extended Figure 6.**
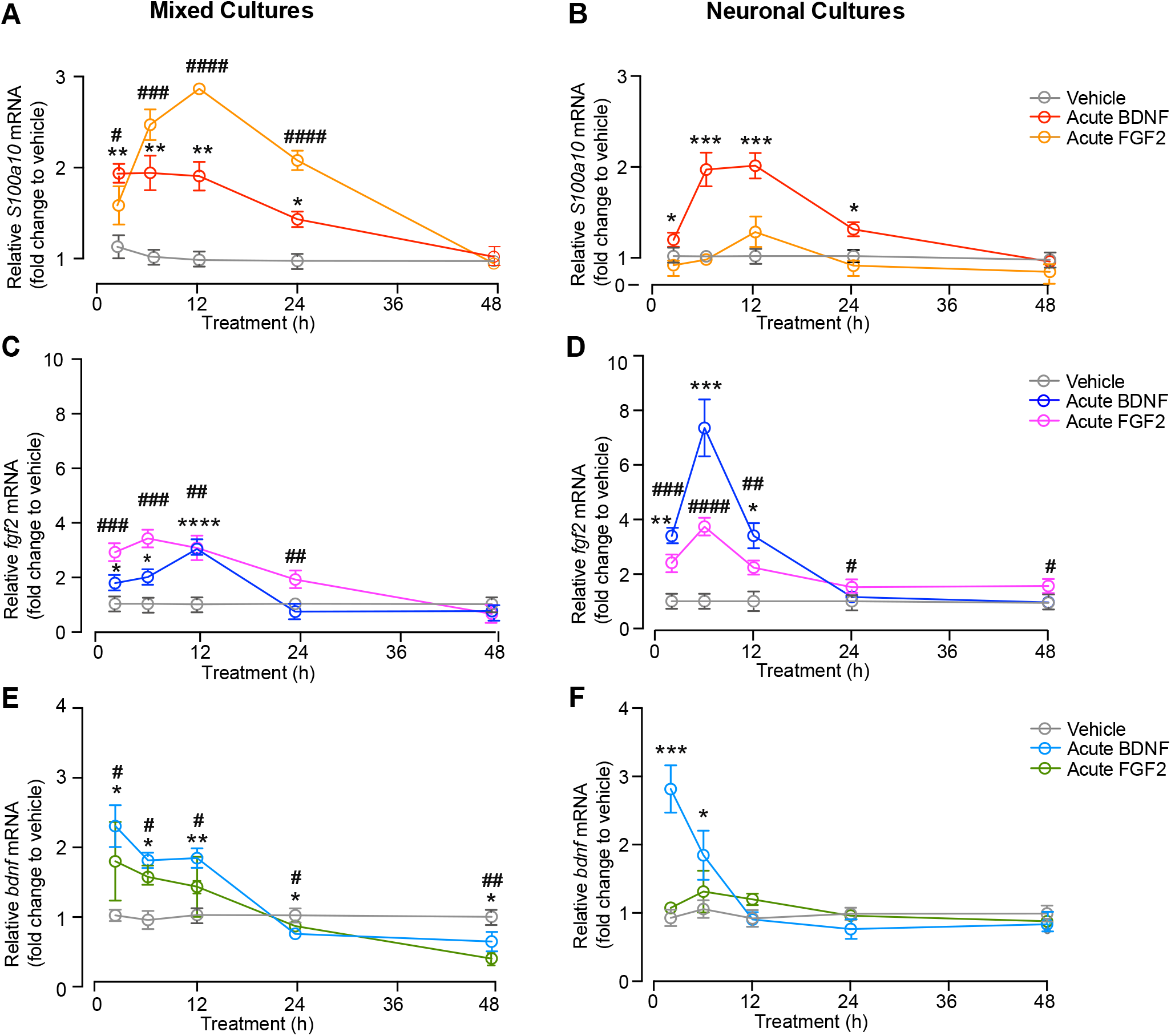
**(A)** Kinetics of *Fgf2, Bdnf* and *S100a10* mRNA expression at 2h, 5h, 12h, 24h, and 48h after acute treatment with FGF2 and BDNF in DIV7 cortical mixed cultures by qRT-PCR. **(B)** Kinetics of *Fgf2, Bdnf* and *S100a10* mRNA expression at 2h, 5h, 12h, 24h, and 48h after acute treatment with FGF2 and BDNF in DIV7 cortical neuronal cultures by qRT-PCR. Statistical analysis was done using one-way ANOVA and corrections for multiple comparisons were performed using post-hoc Bonferroni test. Comparisons were made between vehicle- and FGF2 or BDNF-treated samples; n=6. # symbol indicates comparison between vehicle and FGF2-treated samples, and * symbol indicates comparison between vehicle and BDNF-treated samples. Data are mean +/− SEM; *P≤ 0.05, **P≤ 0.01, ***P≤ 0.005, ****P≤ 0.0005. Experiments were independently reproduced three times with identical results.

