## Supplemental Information for "Neuron-Glia Signaling Regulates the Onset of the Antidepressant Response"

### Extended Table Legends

**Extended Table 1.** Cell populations were annotated based on cell type specific expression of marker genes enriched in each cluster. We obtained 36 distinct clusters in total, which were further grouped and organized into 15 neuronal clusters represented in 15 different sheets. Each sheet represents a cell population depicting marker genes sorted by expression significance based on adjusted p-values (q-value) found using an optimized false discovery rate (FDR) approach.

**Extended Table 2.** Cell populations were annotated based on cell type specific expression of marker genes enriched in each cluster. We obtained 36 distinct clusters in total, which were further grouped and organized into 13 non-neuronal (glial and other) clusters represented in 13 different sheets.

**Extended Table 3.** Cell populations were annotated based on cell type specific expression of marker genes enriched in each cluster. We obtained 36 distinct clusters in total, which were further grouped and organized into 5 inter-neuronal clusters represented in 5 different sheets.

**Extended Table 4.** List of differentially expressed genes for two comparisons: Control versus Stress; Stress versus Stress+Flx treated samples; in the glial, inter-neuronal, and neuronal populations are indicated for three time points of 3, 7, 10 days. Genes that are significantly differentially expressed in both conditions (FDR < 0.1) are shown.

**Extended Table 5.** Gene Ontology enrichment analysis for the list of genes that were affected by stress, then normalized by Flx for three time points of 3, 7, 10 days. Genes that were significantly differentially expressed in both comparisons shown in **Extended Table 4** were identified and Gene Ontology enrichment analysis was performed.

### Extended Materials and Methods

**Mouse primary mixed and neuronal cultures and immunofluorescence analysis.** Pregnant C57Bl/6 mice were purchased from Charles River. E18 Embryos were euthanized, and their cortices isolated. Cortices were isolated from E18 embryos of Pregnant C57Bl/6 mice (Charles River) as described above and a single cell suspension using 0.15% Trypsin-EDTA (Gibco, cat #. 25200056) is made, followed by mechanical trituration. Cells were then plated in Poly-D-Lysine coated coverslips at ~2000 cells/mm (NeuVitro, cat #. GG-18-15-PDL). Cells were cultured and maintained in the respective media for mixed and neuronal cultures as described above. Coverslips were fixed in chilled 4% PFA for 30mins, blocked for 1 hour in 5% Normal Donkey Serum in PBS+0.02% Triton-X, and incubated overnight with primary antibody: GFAP (Abcam, AB4674, 1:1000), Pan-Neuro (MilliporeSigma, MAB2300, 1:500), IBA1 (Wako Chemical, 019-19741, 1:1000), and OLIG2 (R&D Systems, AF2418, 1:1000). Coverslips were then incubated with secondary antibodies: CF488 (SigmaAldrich, SAB4600031, 1:1000), CF555 (SigmaAldrich, SAB4600060, 1:1000), CF594 (SigmaAldrich, SAB4600099, 1:1000), and Alexa 647 (Thermo Fisher Scientific A-21447, 1:500) for 2 hours. Finally, coverslips were stained with DAPI and mounted in ProLong

Gold mountant (Invitrogen, P36930). Slides were imaged on a Zeiss LSM780 AxioObserver confocal microscope equipped with a Plan-Apochromat 20x/0.8 M27 lens. Cell counts were obtained using ImageJ. Briefly, the particle analysis feature of ImageJ was used to count and outline nuclei. Nuclei were then assigned to one of the four cell types based on the intensity of fluorescent signal from each cell marker staining. Cells were counted from n=3 sections, and n=3 replicates and classified per data point.
